# CATH functional families predict protein functional sites

**DOI:** 10.1101/2020.03.23.003012

**Authors:** Sayoni Das, Harry M. Scholes, Christine A. Orengo

## Abstract

**Motivation:** Identification of functional sites in proteins is essential for functional characterisation, variant interpretation and drug design. Several methods are available for predicting either a generic functional site, or specific types of functional site. Here, we present FunSite, a machine learning predictor that identifies catalytic, ligand-binding and protein-protein interaction functional sites using features derived from protein sequence and structure, and evolutionary data from CATH functional families (FunFams).

**Results:** FunSite’s prediction performance was rigorously benchmarked using cross-validation and a holdout dataset. FunSite outperformed all publicly-available functional site prediction methods. We show that conserved residues in FunFams are enriched in functional sites. We found FunSite’s performance depends greatly on the quality of functional site annotations and the information content of FunFams in the training data. Finally, we analyse which structural and evolutionary features are most predictive for functional sites.

**Availability:** The datasets and prediction models are available on request.

**Supplementary information:** Supplementary data are available at *Bioinformatics* online.

## 1 Introduction

Protein functional sites are amino acid residues, or groups of residues, that perform functional roles in proteins. Examples of functional sites include catalytic sites in enzymes, ligand-binding sites for small molecules, metal ions, nucleic acids and other proteins, and protein-protein interaction sites. Characterisation of functional sites is crucial for understanding the molecular basis of life, interpreting the impact of mutations, guiding targeted experiments, protein-engineering and drug design. Computational approaches to functional site prediction is essential since < 1% of all known proteins have any experimentally-characterised or curator-assigned functional site information (UniProtKB, Jan 2019).

A number of different computational strategies have been used to predict generic functional sites (Ashkenazy et al., 2016; Wilkins, Erdin, Lua, & Lichtarge, 2012) and specific types of functional sites, such as catalytic sites (Choudhary, Kumar, Bachhawat, & Pandit, 2017; Wallace, Borkakoti, & Thornton, 1997), ligand-binding sites (Capra, Laskowski, Thornton, Singh, & Funkhouser, 2009; Skolnick & Brylinski, 2009; Wass, Kelley, & Sternberg, 2010) and protein-protein interaction sites (Aumentado-Armstrong, Istrate, & Murgita, 2015; Xue, Dobbs, Bonvin, & Honavar, 2015). Most recent prediction methods use machine learning to predict a specific type of functional site and use features derived from protein sequence and structure (Brylinski & Feinstein, 2013; H. Chen & Zhou, 2005; Zhang et al., 2008). There are considerable differences in the properties of different types of functional sites which provide a basis for these prediction methods. For example, catalytic sites generally have limited exposure to solvent while ligand-binding and protein-binding sites have comparatively higher solvent accessibility. Both catalytic sites and ligand-binding sites are generally well conserved and found in, or near, pockets in the protein structure (Bartlett, Porter, Borkakoti, & Thornton, 2002; Capra et al., 2009). On the other hand, protein interfaces are relatively flat and difficult to predict from sequence conservation alone (Caffrey, Somaroo, Hughes, Mintseris, & Huang, 2004; S. Jones & Thornton, 1997), however, the location of interfaces are conserved (Xue et al., 2015).

The key challenge for predicting functional sites using machine learning is to identify general properties of sites that distinguish one type of site from the others. The features most frequently used by machine learning predictors include sequence conservation information, physico-chemical properties of amino acids, solvent accessibility, secondary structure, pocket information and crystallographic B-factors (Aumentado-Armstrong et al., 2015; Brylinski & Feinstein, 2013; Sankararaman, Sha, Kirsch, Jordan, & Sjölander, 2010; Sun, Wang, Xiong, Hu, & Liu, 2016; Xue et al., 2015). Sequence conservation information is captured using scores derived from multiple-sequence alignments (MSAs) of homologous proteins––based on the assumption that homologous proteins are likely to share functions and, therefore, have similar functional sites. However, this is a non-trivial task, as homologous proteins with similar functions can have diverse functional sites (Brown & Babbitt, 2014; Dessailly, Dawson, Mizuguchi, & Orengo, 2013; Furnham, Dawson, Rahman, Thornton, & Orengo, 2016; Taylor Ringia et al., 2004).

Most functional site predictors use PSI-BLAST (Altschul et al., 1997), which can detect distant homologs, to obtain sequence conservation information. Sequence conservation features can be calculated by an entropy analysis on the resulting position-specific scoring matrix (PSSM), which may include knowledge of positions that are conserved in distantly related proteins. Whilst sequence conservation features calculated this way have been found to predict catalytic and ligand-binding residues well, they are not predictive of protein-protein interfaces (Aumentado-Armstrong et al., 2015).

One way to overcome this is to calculate conservation and other features from protein families where the family grouping brings together sequences that are likely to have similar functional sites. A large number of computational methods utilize protein family information to predict functions or functional sites (Ashkenazy, Erez, Martz, Pupko, & Ben-Tal, 2010; Capra et al., 2009; del Sol Mesa, Pazos, & Valencia, 2003; Innis, Anand, & Sowdhamini, 2004; P. Jones et al., 2014; Lichtarge, Bourne, & Cohen, 1996). However, the quality of protein families greatly affects the performance of prediction, therefore using functionally coherent protein families is crucial.

Functional families (or FunFams) are subfamilies of evolutionary superfamilies in CATH (Sillitoe et al., 2019) which have been generated by a functional classification protocol that is based on recognizing sequence patterns that are conserved in the FunFam but differ between FunFams and, in particular, sequence patterns that reflect specificity determining positions (Das et al., 2016). FunFam relatives have been found to be more functionally similar than other domain-based resources (**see Supplementary Section S1**; (Das et al., 2016). Function prediction pipelines based on FunFams have been consistently ranked among the best function prediction methods by the international CAFA competition (Jiang et al., 2016; Radivojac et al., 2013) and more recently have been amongst the top 5 best performing methods (CAFA3; (Zhou et al., 2019)).

Patterns of conserved residues in FunFam alignments can be used to identify functionally important residues. In fact, conserved residues in FunFam alignments have been found to be highly enriched in known functional sites, including catalytic, ligand-binding, protein interfaces, nucleic acid-binding sites and allosteric sites (see **Supplementary Section S1**; (Das et al., 2016)). However, conserved residues may also include residues important for the folding and packing of the protein. These tend to be conserved across the whole superfamily whereas functional determinants (also known as specificity-determining positions) tend to be differentially conserved in structurally equivalent positions between FunFams, thereby providing insights into the molecular mechanisms of functionally distinct residues in the superfamily (Das et al., 2016; Lee et al., 2016). Therefore, in principle, the finer classification of functional relatives in FunFams, compared to more generic PSI-BLAST approaches, could aid in detecting specific conservation features. These features, combined with other sequence and structure-based features, could be used in an integrated pipeline to predict different types of protein functional sites.

Despite the progress and availability of a wide range of functional site prediction methods, to our knowledge, none of the existing studies have systematically compared the characteristics of the different types of functional sites or attempted to predict multiple functional sites for a given query protein. Moreover, a large number of the prediction methods are not easily accessible or easy to interpret.

In this article, we present a new method (FunSite predictor) for predicting functional sites by using information from a protein functional family classification to predict three types of functional sites: catalytic, ligandbinding and protein-protein interaction sites. This method makes use of features derived from protein sequence, structure and CATH FunFams. Whilst we used the same set of features and machine learning model to train predictors for the three types of functional sites, we used different training sets for each type of site. FunSite predictors outperform all other available functional site predictors. Our combined approach allows the detection of sites which may have multiple functional roles. The performance of the predictors is highly dependent on the quality of the functional site data used for training.

## 2 Methods

### 2.1 Datasets

Three types of functional sites were considered: catalytic sites (CS), ligand-binding sites (LIG) and protein-protein interaction (PPI) sites.

#### 2.1.1 Dataset generation

The general criteria used to generate all the datasets are described in **Supplementary Methods (Section S2).**

##### 2.1.2.1 Catalytic Site (CS) domain datasets

A dataset of 667 domains was generated for catalytic sites from Mechanism and Catalytic Site Atlas (M-CSA; (Ribeiro et al., 2017). Only ‘literature’ type entries were used. 102 of the domains could be mapped to two datasets - the EF_Family dataset (Youn, Peters, Radivojac, & Mooney, 2007) and T124 dataset (Zhang et al., 2008)– previously used to test other predictors (Sun et al., 2016; Zhang et al., 2008). These 102 domains were used to construct a holdout dataset for benchmarking. The remaining 565 domains were used as the training set.

##### 2.1.2.2 Ligand-Binding (LIG) domain datasets

A dataset of 2826 domains was generated for ligand-binding sites for small-molecules (including metals) from BioLip (Yang, Roy, & Zhang, 2013). Ligand-binding site predictions generated by ConCavity (Capra et al., 2009) were available for 800 domains in this dataset. These 800 domains were used to construct a holdout dataset for benchmarking against ConCavity (Capra et al., 2009) and the remaining 2026 domains were used for training.

Subsets of the LIG training and holdout datasets, comprising 675 and 240 domains respectively, gave training and benchmark datasets that only contained metal-binding sites.

##### 2.1.2.3 Protein-Protein Interface (PPI) domain datasets

A dataset of 2247 domains was generated for interchain PPI sites from Inferred Biomolecular Interactions Server (IBIS) (Shoemaker et al., 2012). Only annotations present in experimentally-determined structures were extracted from IBIS. No inferred annotations were used. PPI site predictions were extracted from the meta-PPISP (Qin & Zhou, 2007) server, a meta-predictor of multiple methods, for 600 randomly selected domains in our constructed dataset, of which 599 domains returned results. These 599 domains were used as the holdout dataset for benchmarking against meta-PPISP.

##### 2.1.2.4 Combined dataset of CS, LIG and PPI domains

A dataset of 175 enzyme domains was generated from the CS, LIG and PPI datasets that have at least one annotation in each domain, for all the three types of sites.

More information about the datasets is provided in Supplementary Table S1. As the number of functional site residues in a protein domain is generally much lower than the number of non-site residues, the datasets have large class imbalance discussed further below together with strategies for addressing this. (**Supplementary Figure S1**).

### 2.2 FunSite predictors

#### 2.2.1 Implementation and Training

To build the FunSite predictors we used gradient boosted decision trees (Friedman, 2001), applying the XGBoost (eXtreme Gradient Boosting) implementation (T. Chen & Guestrin, 2016). Gradient boosted trees are ensembles of weakly learning decision trees. During training, the XGBoost algorithm generates an ordered ensemble of decision trees in serial that are each fit to the residual error generated by all preceding trees in the ensemble. As such, gradient boosted trees are able to achieve high performance using only weakly learning decision trees.

The scikit-learn (Pedregosa et al., 2011) library in Python (www.python.org) was used to train and evaluate predictors using XGBoost’s scikit-learn API. The number of decision trees in the ensemble was set to 1000. XGBoost hyperparameters were optimised using grid search with 5-fold cross validation (GridSearchCV function in scikit-learn). The hyperparameters and hyperparameter ranges used for the different FunSite predictors are listed in **Supplementary Table S2**. To ensure the prediction error was not underestimated, the GroupKfold function in scikit-learn was used to select non-overlapping fold groups during cross-validation, such that residues from the same domain do not appear in the sets used for training and testing.

The ratio of site residues (positive samples) to non-site residues (negative samples) was very low for CS (1:63) and LIG (1:23) datasets (see **Supplementary Table S1, Supplementary Figure S2**). In order to reduce the class imbalance during training, the site to non-site ratio for the CS and LIG predictors was set to 1:6 by randomly selecting non-site residues. This is similar to class distributions used in previous studies (Sun et al., 2016). No changes were made to the non-site set for PPI as the ratio of site to non-site residues in the PPI training set was 1:4.

#### 2.2.2 Feature generation

For each residue, FunSite generates features based on sequence, structure and FunFams. The sequence- and structure-based features generated are listed in **Supplementary Methods (Section S2)**. The novel FunFam-based protein family features are described below:

##### 2.2.2.1 Protein family features

To generate protein family features, each query protein sequence was scanned against the library of CATH FunFam HMM models using HMMER3 (Eddy, 2010). cath-resolve-hits (Lewis, Sillitoe, & Lees, 2018) was used to determine the optimal assignment of the sequence into domain regions based on the HMM matches. Regions of the query sequence were assigned to a FunFam if the HMM match E-value was smaller than the model inclusion threshold, defined as the maximum E-value obtained by scanning each sequence within a particular FunFam against its HMM. The query sequence was then aligned with the sequences in the FunFam alignment using the mafft-add option in MAFFT (Katoh & Standley, 2013). Following alignment of domain sequences to CATH FunFams the following features were calculated:

i. *Evolutionary conservation scores:* The evolutionary conservation score for each residue was calculated from the FunFam multiple sequence alignment using Scorecons (Valdar, 2002). This is only possible for FunFams with sufficient information content in their alignment (see **Supplementary Section S1**). Scorecons scores range from 0 to 1 and residues having scores ≥ 0.7 are considered to be highly conserved (Das et al., 2016).
ii. *PSSM and weighted observed percentages (WOP) features:* PSSM and weighted observed percentages (WOP) were calculated for the FunFam alignment using PSI-BLAST. 20-dimensional PSSM and 20-dimensional WOP vectors are obtained for each residue.
iii. *Conservation scores and predicted functional determinant (FD) scores from Structural Clusters of FunFams:* For each CATH superfamily, FunFam alignments with at least one domain structure, were merged to form Structural Clusters (SC5), such that structural relatives from the merged FunFams can be superimposed within a 5Å RMSD threshold. The Structural Cluster (SC5) MSAs were generated by merging FunFam MSAs guided by the structural alignment of domain structures from each constituent FunFam. Conservation scores for SC5 alignments were calculated by Scorecons. Functional determinant (FD) scores were predicted using GroupSim (Capra & Singh, 2008). GroupSim scores range from 0 to 1 where a higher score indicates higher probability for a column in an alignment to be a FD (i.e. to be associated with different conservation characteristics between FunFams within the structural cluster).

#### 2.2.3 FunSite prediction protocol

The FunSite prediction protocol is shown in **Figure 1**. For each query protein, the residue feature vector (**see Section 2.2.2**) used by the FunSite predictors was generated for all residues. The three FunSite predictors (FunSite-CS, FunSite-LIG and FunSite-PPI) were run separately on the residue vectors of each query protein to generate separate prediction scores for each residue to be a catalytic, ligand-binding or protein-protein interaction site residue. The three predictors were run separately, instead of combining them into a single model, because the functional plasticity of residues (Dessailly et al., 2013) means that, depending on its context, a residue can perform multiple functions. For example, some proteins often use the same or overlapping set of surface residues to bind small molecules and other proteins (Davis & Sali, 2010; Mohamed, Degac, & Helms, 2015). Moreover, an increasing number of proteins have been found to perform multiple molecular functions using different, overlapping or sometimes the same set of residues (Das, Khan, Kihara, & Orengo, 2017). Therefore, the aim is to predict probabilities for all three types of site. Furthermore, to remove false positives, residues that are positively predicted to be a particular functional type are filtered out if there are no other positively predicted site residues of the same type in their structural neighborhood (5Å). This is based on the assumption that functional sites generally comprise two or more residues. This filtering step is applied when making combined predictions (i.e. all types of functional sites) but not for the individual predictors.

**Figure 1.**
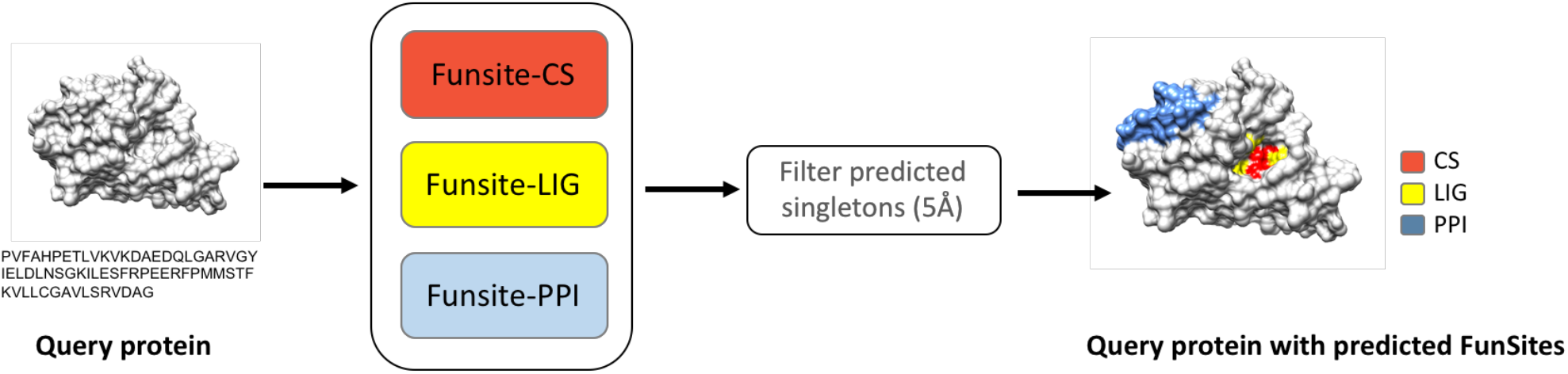
FunSite prediction method.

### 2.3 Comparison with other predictors

Performance of the FunSite predictors was benchmarked against the following functional site prediction methods:

#### 2.3.1 Publicly available predictors

The following prediction methods were used, available at the time of writing.

*Catalytic site predictors:* CSmetaPred was used for benchmarking the predicted CS sites by FunSite-CS predictor. This is a consensus meta-predictor that also provides predictions for its three component predictors– CRpred (Zhang et al., 2008), DISCERN (Sankararaman et al., 2010) and EXIA2 (Lu, Yu, Chien, & Huang, 2014), all of which exploit machine learning.

*Ligand-binding site predictor:* ConCavity (Capra et al., 2009) was used for benchmarking the predicted LIG sites by FunSite-LIG as its predictions were easily available.

*Protein-protein interaction site predictors:* The meta-predictor meta-PPISP (Qin & Zhou, 2007) was used as this does not require partner-specific information for generating predictions. meta-PPISP also provides the predictions for three component independent machine learning predictors––PINUP (Liang, Zhang, Liu, & Zhou, 2006), Promate (Neuvirth, Raz, & Schreiber, 2004) and cons-PPISP (Chen and Zhou 2005).

#### 2.3.2 Generic predictors

Generic predictors were also constructed for benchmarking the performance of the FunSite predictors. This was done by using the same training and test datasets, model implementation and parameters for model training. However, the generic predictors did not include any FunFam-based features. Instead, they include similar conservation-based features derived from PSSMs generated by PSI-BLAST. These features were combined with the other sequence- and structure-based features used by FunSite and which are frequently used by other functional site predictors, such as amino acid properties, pocket predictions, solvent accessibility, secondary structure information and B-factor. The full list of features used for training the generic predictors is provided in **Supplementary Table S3**.

The generic predictors provide an unbiased benchmark that is not affected by the size of the training dataset, the sampling strategy used, the machine learning model, or their different implementations. As a result, any improvement of performance shown by a FunSite predictor over its generic equivalent can be assumed to be solely due to the power of the FunFam-based features.

The evaluation methods used to compare performance with other predictors are described in **Supplementary Methods (Section S2).**

## 3 Results and Discussion

### 3.1 Analysis of conservation properties for different functional site types

Residue conservation analysis of CATH FunFam alignments with high information content showed that experimentally characterised catalytic and ligand-binding site residues are generally highly conserved, i.e. they have Scorecons conservation scores ≥ 0.7. Figure 2 shows a comparison of the distribution of conservation scores for different types of known functional site residues identified in CATH FunFams and non-site residues that are either buried or present on the protein surface. The majority of known catalytic site residues correspond to highly conserved residues in CATH FunFams. In a few cases, catalytic residues have low conservation scores due to the presence of mutated residues in those sites in some of the domains. A large proportion of the known ligand-binding sites were also found to be highly conserved in CATH FunFams. Comparatively, only a small number of known protein-protein interaction sites were found to be conserved in CATH FunFams. This is consistent with the findings of previous studies suggesting that few residues are conserved in protein interfaces (Caffrey et al., 2004; David & Sternberg, 2015; Humphris & Kortemme, 2007).

**Figure 2.**
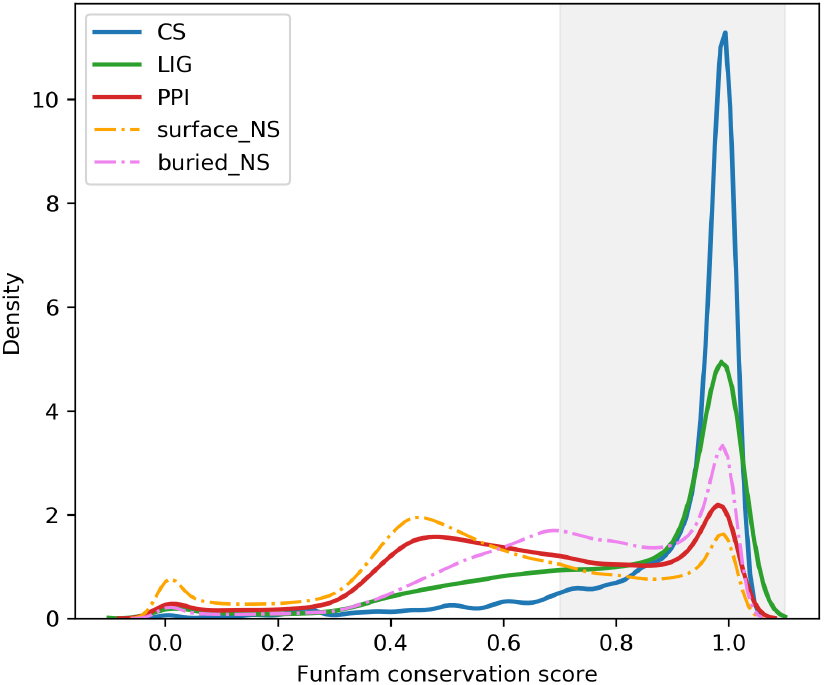
Density plots showing the distribution of sequence conservation scores for catalytic site (CS), ligand-binding site (LIG) and protein-protein interaction site (PPI) residues in CATH FunFams compared to non-site residues that are buried (buried_NS) and those that are on the surface (surface_NS). The sequence conservation scores were generated by Scorecons (ranging from 0-1) and the residues with scores ≥ 0.7 are considered to be conserved (shaded gray in the figure).

Buried non-site residues were also observed to be under evolutionary constraints because of their role in maintaining protein folding and stability. Non-site residues present on the surface were found to be the least conserved, although it is important to note that some of these may include sites that have not yet been annotated and other types of functional sites not considered in this analysis, such as allosteric and phosphorylation sites.

### 3.2 Selection of discriminating FunSite features for sites

Knowledge of the general characteristics of functional site residues was used to select a feature set for training the predictors to distinguish functional site residues from non-functional site residues. Some of these features included traditional characteristics such as sequence conservation scores **(see Supplementary Figure S2a)**, solvent accessibility values and predicted pocket residues that are known to capture discriminating characteristics of CS, LIG and PPI residues. Additionally, a large number of other features were included in the FunSite predictors that were derived from combining sequence conservation information with other structural characteristics. For example, the number of residues surrounding conserved pocket residues (i.e. at a distance < 5Å) was found to be significantly different between site and non-site residues for CS and LIG **(see Supplementary Figure S2b)** and the average number of surface residues within a 5Å radius **(see Supplementary Figure S2c)** was found to differentiate PPI residues from other residues.

### 3.3 Performance evaluation of FunSite predictors

The performance of the FunSite predictors for CS, LIG and PPI sites was evaluated and compared to generic predictors and other publicly available functional site predictors.

#### 3.3.1 FunSite-CS

The precision-recall (PR) curves in **Figure 3a** show the performance of FunSite-CS in predicting catalytic sites, compared to the generic predictor. Both FunSite and generic predictors were generated from 5-fold cross-validation of the CS training dataset. FunSite-CS performs better with respect to both precision and recall and gives a higher area under the PR curve (PR AUC). Other evaluation metrics calculated on the CS training set are shown in **Supplementary Table S4**.

**Figure 3.**
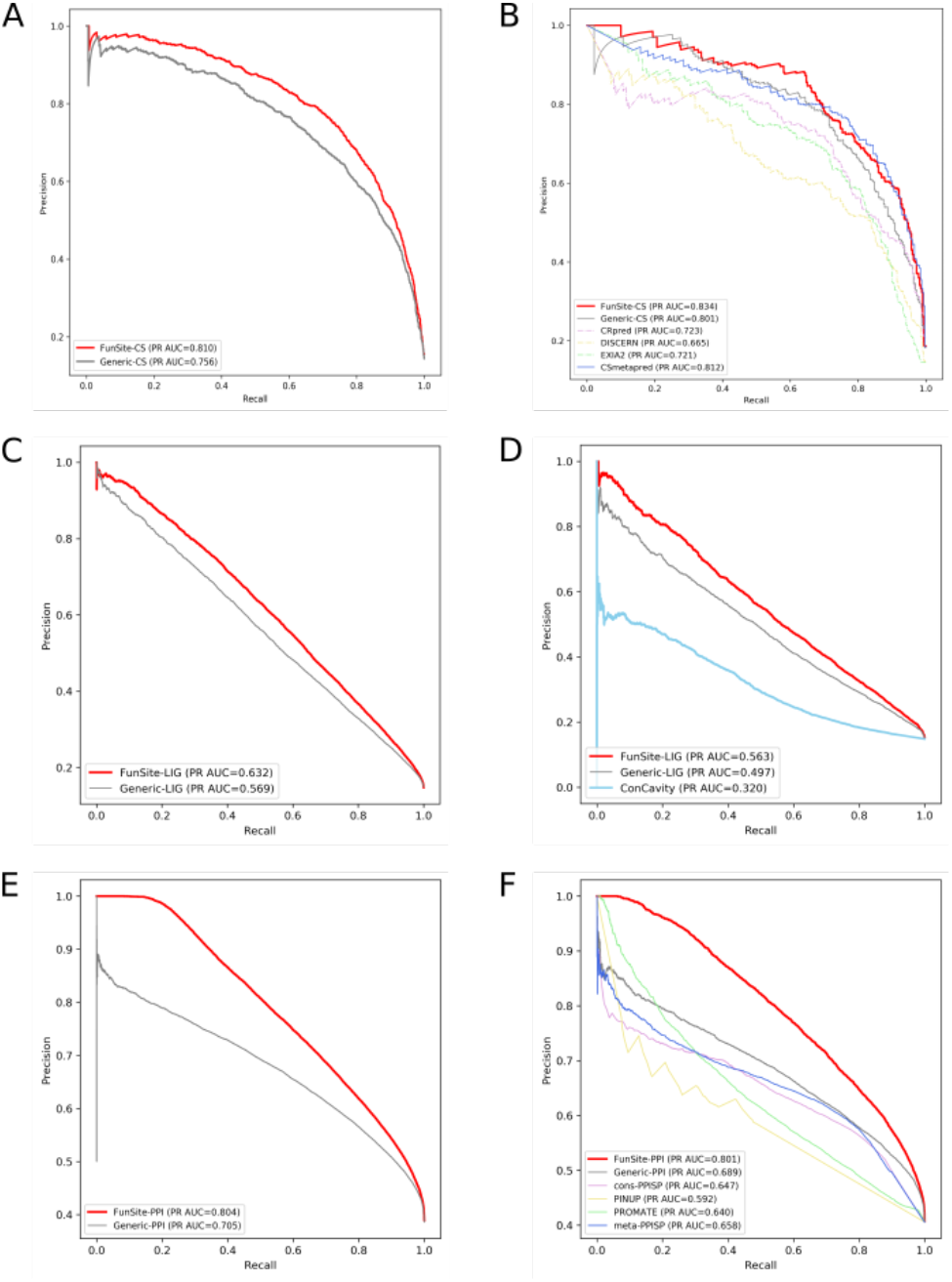
Precision-recall curves showing the performance of the FunSite predictors for catalytic (CS), ligand-binding (LIG) and protein-protein interaction (PPI) sites. Figures (a), (c) and (e) show the 5-fold cross validation results on the training datasets for the FunSite-CS, FunSite-LIG and FunSite-PPI predictors respectively compared to the generic predictors. Figures (b), (d) and (f) show the performance of the FunSite-CS, FunSite-LIG and FunSite-PPI predictor on the hold-out test sets compared to other predictors.

**Figure 3b** shows the performance of FunSite-CS compared to other predictors on the CS holdout set of 106 domains. The FunSite-CS predictor performs competitively with the meta-predictor CSmetapred and performs better than the generic CS predictor and the individual catalytic site predictors (CRpred, EXIA2 and DISCERN). The distribution of prediction probabilities by FunSite-CS predictor on the holdout set is shown in **Supplementary Figure S3a**. It is important to note that the generic predictor also outperforms the publicly available predictors. This is most likely due to the use of a larger training set for the generic predictor and increase in the quality of catalytic site annotations (Ribeiro et al. 2017).

#### 3.3.2 FunSite-LIG

**Figure 3c** shows the performance of the FunSite-LIG predictor from 5-fold cross-validation of the LIG training set compared to the generic LIG predictor. Neither predictors perform well, as shown by the shape and area under the precision-recall curves. However, when the LIGmetal dataset (a subset of the LIG dataset that only contains metal-binding sites) was used in a similar manner, to predict metal-binding sites, a large improvement was observed on the performance of both the FunSite-LIGmetal and generic LIGmetal predictors (**Supplementary Figure S4**). Other evaluation metrics calculated on the LIG test dataset are shown in **Supplementary table S4.**

The comparatively lower performance of the predictors for the LIG set is probably due to variations in the characteristics of the ligand binding set. For example, this could include (i) differences in the types of ligands represented in the dataset (e.g. cognate and non-cognate) and (ii) differences in the characteristics of ligand binding between enzymes and non-enzymatic proteins. For example, it is known that non-cognate ligands can bind to different regions in a protein to those bound by the cognate ligands, reflecting different binding characteristics (Tyzack, Fernando, Ribeiro, Borkakoti, & Thornton, 2018).

**Figure 3d** shows the performance of the LIG predictors on the LIG hold-out test set compared to the ConCavity ligand-binding site prediction method. The FunSite-LIG predictor shows the highest PR AUC, followed by the generic LIG predictor and then ConCavity. The FunSite-LIGmetal predictor also performs better than the generic predictor on the LIGmetal test set (**Supplementary Figure S4**).

#### 3.3.3 FunSite-PPI

**Figure 3e** shows the performance of the FunSite-PPI and generic-PPI predictors derived from 5-fold cross-validation of the PPI training dataset. FunSite-PPI outperforms the generic predictor to a greater extent than both FunSite-CS and FunSite-LIG. This shows that the specific conservation-based features of the FunSite-PPI predictor make a more powerful contribution towards distinguishing between protein interfaces and other residues, than the conservation measures captured from the PSI-BLAST PSSM.

On the PPI holdout set, the FunSite predictor also outperformed the generic predictor and the meta-predictor meta-PPISP along with the individual PPI predictors (cons-PPISP, PINUP and PROMATE) **(Figure 3f).** Other evaluation metrics calculated on the PPI test dataset are shown in **Supplementary table S4**.

### 3.4 Most influential features for FunSite predictions

The most important features for the FunSite predictions were analysed using Tree SHAP (Lundberg & Lee, 2017). SHAP combines a number of approaches to identify the influence of individual features on the prediction performance of tree ensemble methods. Feature influences are reported as SHAP values which are more consistent than classic feature importance metrics (Lundberg, Erion, & Lee, 2018). Higher SHAP values for features correspond to higher log-odds ratios that a residue is a functional site. **Figure 4** shows the top 25 features for the FunSite-CS, -LIG and -PPI predictors ranked by SHAP values (see **Supplementary Figure S5** for FunSite-LIGmetal). The top features are conservation scores, hydration potential, pocket information for CS sites; conservation scores, pocket information, solvent accessibility for LIG sites; and solvent accessibility, residue curvature for PPI sites.

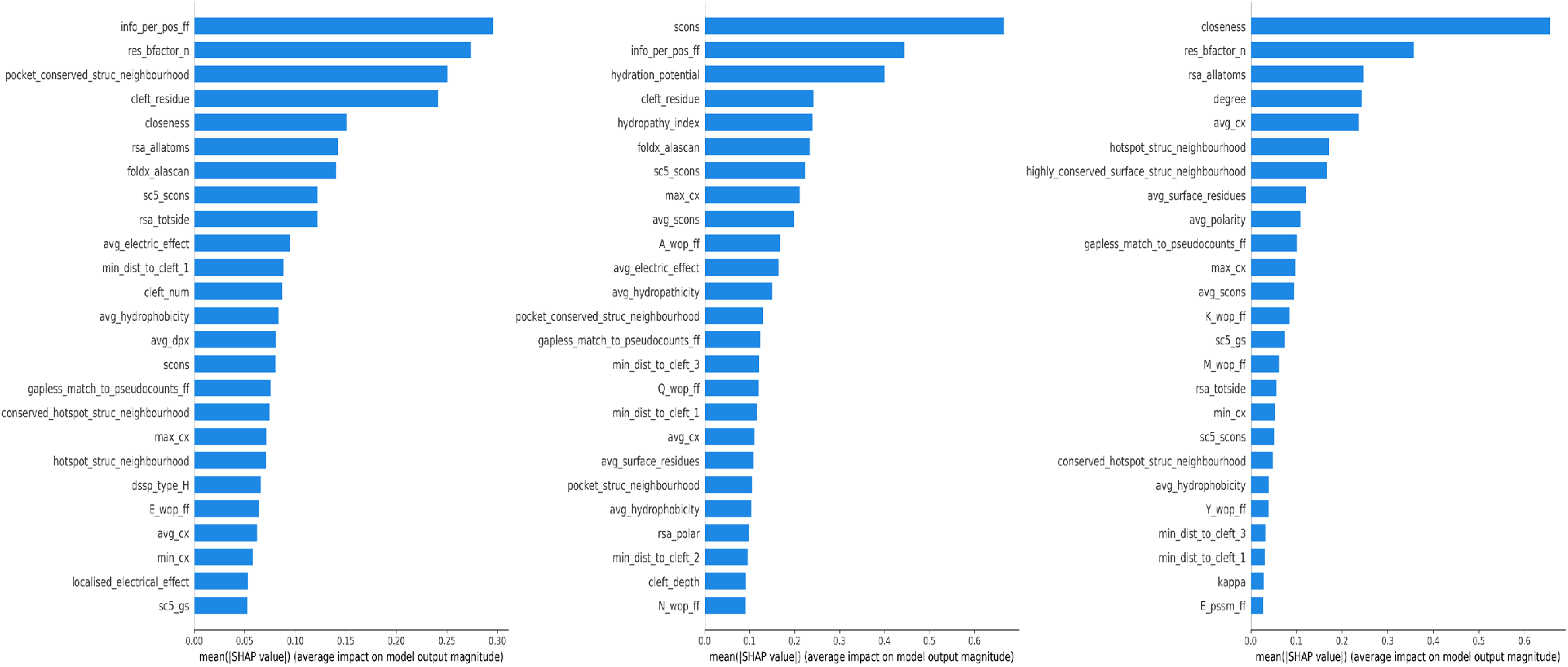

### 3.5 FunSite predictions for the combined dataset

The full FunSite prediction protocol (**Figure 1**) was run on the combined dataset of 175 enzyme domains that have CSA, LIG and PPI functional site annotations. Prediction scores for each FunSite predictor were generated for all residues. To do this the three separate FunSite predictors were independently trained on the CS, LIG and PPI training datasets, respectively, from which the 175 query domains had been removed. As in the previous tests, positively predicted residues that had no other surrounding predicted residue of the same type were filtered out.

For each domain in the dataset, residues were assigned three ranks, based on the prediction scores from each of the three FunSite predictors. Only positively predicted residues, with predicted probability ≥ 0.5, for each site were ranked. The top ranked residues, up to a maximum of 20 residues, for each type of functional site were generated as FunSite predictions. As an example, **Figure 5** shows the FunSite predictions obtained for the12asA00 domain.

**Figure 5.**
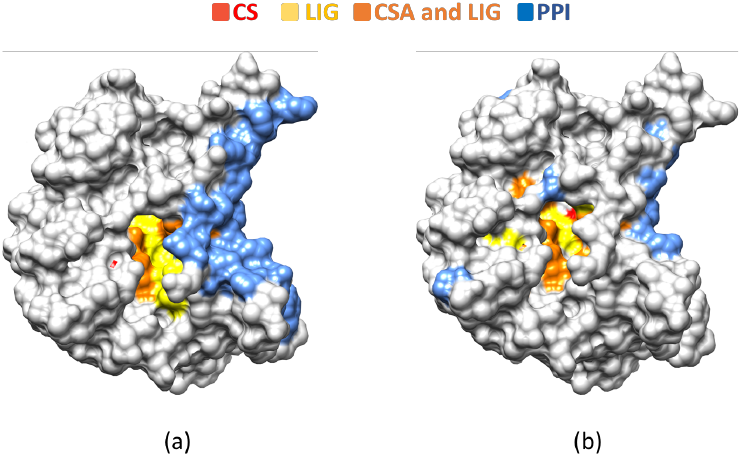
Example domain (12asA00) (a) Known functional sites (b) Predicted functional sites

The final FunSite predictions (i.e. top 20 predicted residues for each predictor) were found to be significantly more enriched (**Table 1, Supplementary Figure S6**) in known functional sites compared to FunSite predictions which hadn’t been filtered by removing likely false positives. They were also enriched compared with sets obtained by simply predicting that the conserved sites in FunFams are functional sites. The filtering step was most valuable for removing false positives in the prediction of PPI sites (**Table 1**).

**Table 1.** Results from one-sided Fisher’s exact tests comparing the precision/enrichment of the prediction of functional sites using FunSite predictor with filtering, compared to no filtering, and compared to simply using conserved sites as predicted functional sites.

|  | FunSite-CS | FunSite-LIG | FunSite-PPI |
| --- | --- | --- | --- |
| FunSite vs FunSite_no-filter | 5.6e-05 | 1.4e-09 | 7.7e-175 |
| FunSite vs conserved site | 6e-181 | 0 | 0 |

**Supplementary Figure S7** shows the pairwise plot of the prediction scores for the three different types of predicted sites in the 175 domains of the combined dataset. It can be seen that the prediction scores for CS sites are highly correlated with LIG sites. PPI sites do not show any positive correlation with either CS or LIG sites, although some sites have a high PPI score and either a high CS or high LIG score. This will occur in proteins that have an active site located between two or more domains (Ali, 2005; Der, 2012). There are also cases where LIG sites overlap with PPI sites (Mohamed, Degac, & Helms, 2015).

## Conclusions

We have developed predictors for three types of functional site residues in proteins: catalytic, ligand-binding and protein interface residues. Our prediction methods exploit gradient boosted decision trees and a range of features based on sequence, structure and protein families. Features include conservation scores for residues, measured using entropy-based analysis of the multiple sequence alignment (MSA) of the functional family to which the protein domain has been assigned. Conservation analysis of residues known to be functional sites showed that catalytic sites were associated with high conservation scores. Ligand binding sites also exhibit high conservation scores, although not as high as catalytic residues. Protein interface residues, whilst showing some degree of conservation, exhibit the lowest level of conservation out of all three functional types.

Our method is focused at the domain level since FunFams are classified at the domain level and domains often have a particular functional role that is independent of the multi-domain context (Bashton & Chothia, 2007). In the context of catalytic residues, studies of enzyme families have shown that the majority of catalytic residues typically lie in one of the domains in a multi-domain protein (Furnham et al., 2016). In addition to using conservation features obtained from entropy analysis of FunFam MSAs, we also explored a range of other structure- and sequence-based features typically used in the prediction of protein functional sites. SHAP analysis revealed that conservation scores figured highly for catalytic and ligand binding residues, together with pocket information, whilst accessibility and curvature were more important than conservation for protein interface residues, although conservation was a contributing feature. In fact, we observed that the difference in performance between the FunSite predictor and the generic predictor was greatest for protein interface residues, suggesting that conservation measures derived from the FunFam MSAs were better able to distinguish interface/non-interface than those derived from the generic PSSMs.

A major challenge in assessing the performance of site predictors is the difficulty in obtaining appropriately annotated functional sites. Recent studies of ligand-binding data in the PDB revealed that in many cases the bound ligands were not cognate (Tyzack et al., 2018). In addition, databases can employ different definitions of these sites. This will introduce both false positives (i.e. from non-cognate ligand-binding residues) and false negatives (i.e. missing cognate ligand-binding residues) confounding performance measures. The improvement observed in prediction performance for the metal ligand binding predictor is quite likely to be partially attributable to the greater accuracy of the positive annotations in this case. Higher quality annotations of functional sites will improve the reported performance of prediction methods, as demonstrated by the performance of our generic predictors.

Despite reasonable performances for CSA and LIG prediction, predicting protein interface residues is challenging. This could be aided by including features associated with co-varying residues (De Juan, Pazos, & Valencia, 2013). However, these predictions require knowledge of the protein partner and exploiting that information was outside the scope of this analysis. Future work will explore the enhancements in performance which can be obtained by using this type of information.

The main novelty of our approach lies in the fact that we are able to consider the likelihood of residues in a query protein belonging to three different types of functional site and we allow for the possibility that a residue may have more than one functional role, such as contributing to both the binding of a domain or protein partner and to the binding of the substrate.

Since our approach has the advantage over other publicly available methods of exploiting recent, more comprehensive and possibly more accurate data for training the predictors, we also compared our approach against a generic predictor which uses the same features as our predictor, except for the conserved residue features. These were obtained from multiple sequence alignments obtained by PSI-BLAST rather than FunFams. This allowed us to assess the benefits of deriving conservation data from more functionally pure sets of relatives. We found that our FunFam based predictors outperformed the generic predictor for all types of functional sites.

Future work involves extending our pipeline to other types of functional sites and applying our predictor to identify putative functional sites for all FunFams in CATH superfamilies with high-quality conservation data from high information content from sequence diverse relatives.

## Acknowledgements

The authors would like to thank Dr. Jonathan Lees and Dr. Ian Sillitoe for their helpful advice on various aspects of this work.

## References

Ali, M., & Imperiali, B. (2005). Protein oligomerization: How and why. Bioorganic & Medicinal Chemistry, 13(17) 5013–5020.

Altschul, S. F., Madden, T. L., Schäffer, A. A., Zhang, J., Zhang, Z., Miller, W., & Lipman, D. J. (1997). Gapped {BLAST} and {PSI-BLAST}: a new generation of protein database search programs. Nucleic Acids Res., 25(17), 3389–3402.

Ashkenazy, H., Abadi, S., Martz, E., Chay, O., Mayrose, I., Pupko, T., & Ben-Tal, N. (2016). {ConSurf} 2016: an improved methodology to estimate and visualize evolutionary conservation in macromolecules. Nucleic Acids Res., 44(W1), W344–50.

Ashkenazy, H., Erez, E., Martz, E., Pupko, T., & Ben-Tal, N. (2010). ConSurf 2010: calculating evolutionary conservation in sequence and structure of proteins and nucleic acids. Nucleic Acids Research, 38(suppl_2), W529–W533.

Aumentado-Armstrong, T. T., Istrate, B., & Murgita, R. A. (2015). Algorithmic approaches to protein-protein interaction site prediction. Algorithms Mol. Biol., 10, 7.

Bartlett, G. J., Porter, C. T., Borkakoti, N., & Thornton, J. M. (2002). Analysis of catalytic residues in enzyme active sites. J. Mol. Biol., 324(1), 105–121 https://doi.org/10.1016/S0022-2836(02)01036-7

Bashton, M., & Chothia, C. (2007). The generation of new protein functions by the combination of domains. Structure, 15(1), 85–99.

Brown, S. D., & Babbitt, P. C. (2014). New insights about enzyme evolution from large scale studies of sequence and structure relationships. J. Biol. Chem., 289(44), 30221–30228.

Brylinski, M., & Feinstein, W. P. (2013). {eFindSite}: improved prediction of ligand binding sites in protein models using meta-threading, machine learning and auxiliary ligands. J. Comput. Aided Mol. Des., 27(6), 551–567.

Caffrey, D. R., Somaroo, S., Hughes, J. D., Mintseris, J., & Huang, E. S. (2004). Are protein-protein interfaces more conserved in sequence than the rest of the protein surface? Protein Science, 13(1), 190–202. https://doi.org/10.1110/ps.03323604

Capra, J. A., Laskowski, R. A., Thornton, J. M., Singh, M., & Funkhouser, T. A. (2009). Predicting Protein Ligand Binding Sites by Combining Evolutionary Sequence Conservation and 3D Structure. PLoS Computational Biology, 5(12), e1000585. https://doi.org/10.1371/journal.pcbi.1000585

Capra, J. A., & Singh, M. (2008). Characterization and prediction of residues determining protein functional specificity. Bioinformatics, 24(13), 1473–1480.

Chen, H., & Zhou, H.-X. (2005). Prediction of Interface Residues in Protein–Protein Complexes by a Consensus Neural Network Method: Test Against NMR Data. Proteins: Structure, Function, and Bioinformatics, 61(1), 21–35. https://doi.org/10.1002/prot.20514

Chen, T., & Guestrin, C. (2016). {XGBoost}: A Scalable Tree Boosting System. In Proceedings of the 22Nd {ACM} {SIGKDD} International Conference on Knowledge Discovery and Data Mining (pp. 785–794). New York, NY, USA: ACM.

Choudhary, P., Kumar, S., Bachhawat, A. K., & Pandit, S. B. (2017). {CSmetaPred}: a consensus method for prediction of catalytic residues. BMC Bioinformatics, 18(1), 583.

Das, S., Khan, I., Kihara, D., & Orengo, C. (2017). Exploring Structure--Function Relationships in Moonlighting Proteins. In Moonlighting Proteins: Novel Virulence Factors in Bacterial Infections (pp. 21–43). John Wiley & Sons Hoboken.

Das, S., Lee, D., Sillitoe, I., Dawson, N. L., Lees, J. G., & Orengo, C. A. (2016). Functional classification of {CATH} superfamilies: a domain-based approach for protein function annotation. Bioinformatics, 32(18), 2889.

David, A., & Sternberg, M. J. E. (2015). The Contribution of Missense Mutations in Core and Rim Residues of Protein-Protein Interfaces to Human Disease. Journal of Molecular Biology. https://doi.org/10.1016/j.jmb.2015.07.004

Davis, F. P., & Sali, A. (2010). The overlap of small molecule and protein binding sites within families of protein structures. PLoS Computational Biology, 6(2).

De Juan, D., Pazos, F., & Valencia, A. (2013). Emerging methods in protein co-evolution. Nature Reviews Genetics, 14(4), 249–261.

del Sol Mesa, A., Pazos, F., & Valencia, A. (2003). Automatic methods for predicting functionally important residues. Journal of Molecular Biology, 326(4), 1289–1302.

Der, B., Edwards, D., & Kuhlman, B. (2012). Catalysis by a de novo zinc-mediated protein interface: implications for natural enzyme evolution and rational enzyme engineering. Biochemistry, 51(18), 3933–3940.

Dessailly, B. H., Dawson, N. L., Mizuguchi, K., & Orengo, C. A. (2013). Functional site plasticity in domain superfamilies. Biochim. Biophys. Acta, 1834(5), 874–889.

Eddy, S. (2010). {HMMER3}: a new generation of sequence homology search software. URL: http://Hmmer.Janelia.Org.

Friedman, J. H. (2001). Greedy function approximation: a gradient boosting machine. Annals of Statistics, 1189–1232.

Furnham, N., Dawson, N. L., Rahman, S. A., Thornton, J. M., & Orengo, C. A. (2016). {Large-Scale} Analysis Exploring Evolution of Catalytic Machineries and Mechanisms in Enzyme Superfamilies. J. Mol. Biol., 428(2 Pt A), 253–267.

Humphris, E. L., & Kortemme, T. (2007). Design of multi-specificity in protein interfaces. PLoS Computational Biology, 3(8).

Innis, C. A., Anand, A. P., & Sowdhamini, R. (2004). Prediction of functional sites in proteins using conserved functional group analysis. Journal of Molecular Biology, 337(4), 1053–1068.

Jiang, Y., Oron, T. R. T. R., Clark, W. T. W. T., Bankapur, A. R. A. R., D’Andrea, D., Lepore, R., … Radivojac, P. (2016). An expanded evaluation of protein function prediction methods shows an improvement in accuracy. Genome Biology, 17(1), 184. https://doi.org/10.1186/s13059-016-1037-6

Jones, P., Binns, D., Chang, H.-Y., Fraser, M., Li, W., McAnulla, C., … others. (2014). InterProScan 5: genome-scale protein function classification. Bioinformatics, 30(9), 1236–1240.

Jones, S., & Thornton, J. M. (1997). Analysis of protein-protein interaction sites using surface patches. J. Mol. Biol., 272(1), 121–132.

Katoh, K., & Standley, D. M. (2013). {MAFFT} multiple sequence alignment software version 7: improvements in performance and usability. Mol. Biol. Evol., 30(4), 772–780.

Lee, D., Das, S., Dawson, N. L., Dobrijevic, D., Ward, J., & Orengo, C. (2016). Novel Computational Protocols for Functionally Classifying and Characterising Serine {Beta-Lactamases}. PLoS Comput. Biol., 12(6), e1004926.

Lewis, T. E., Sillitoe, I., & Lees, J. G. (2018). cath-resolve-hits: a new tool that resolves domain matches suspiciously quickly. Bioinformatics.

Liang, S., Zhang, C., Liu, S., & Zhou, Y. (2006). Protein binding site prediction using an empirical scoring function. Nucleic Acids Res., 34(13), 3698–3707. https://doi.org/10.1093/nar/gkl454

Lichtarge, O., Bourne, H. R., & Cohen, F. E. (1996). An evolutionary trace method defines binding surfaces common to protein families. Journal of Molecular Biology, 257(2), 342–358.

Lu, C.-H., Yu, C.-S., Chien, Y.-T., & Huang, S.-W. (2014). EXIA2: web server of accurate and rapid protein catalytic residue prediction. BioMed Research International, 2014, 807839. https://doi.org/10.1155/2014/807839

Lundberg, S. M., Erion, G. G., & Lee, S.-I. (2018). Consistent Individualized Feature Attribution for Tree Ensembles.

Lundberg, S. M., & Lee, S.-I. (2017). A Unified Approach to Interpreting Model Predictions. In I. Guyon, U. V Luxburg, S. Bengio, H. Wallach, R. Fergus, S. Vishwanathan, & R. Garnett (Eds.), Advances in Neural Information Processing Systems 30 (pp. 4765–4774). Curran Associates, Inc.

Mohamed, R., Degac, J., & Helms, V. (2015). Composition of overlapping protein-protein and protein-ligand interfaces. PloS One, 10(10).

Neuvirth, H., Raz, R., & Schreiber, G. (2004). ProMate: A Structure Based Prediction Program to Identify the Location of Protein–Protein Binding Sites. Journal of Molecular Biology, 338(1), 181–199. https://doi.org/10.1016/J.JMB.2004.02.040

Pedregosa, F., Varoquaux, G., Gramfort, A., Michel, V., Thirion, B., Grisel, O., … Duchesnay, É. (2011). Scikit-learn: Machine Learning in Python. J. Mach. Learn. Res., 12(Oct), 2825–2830.

Qin, S., & Zhou, H.-X. (2007). meta-PPISP: a meta web server for protein-protein interaction site prediction. Bioinformatics, 23(24), 3386–3387. https://doi.org/10.1093/bioinformatics/btm434

Radivojac, P., Clark, W. T., Oron, T. R., Schnoes, A. M., Wittkop, T., Sokolov, A., … Friedberg, I. (2013). A large-scale evaluation of computational protein function prediction. Nature Methods, 10(3), 221–227. https://doi.org/10.1038/nmeth.2340

Ribeiro, A. J. M., Holliday, G. L., Furnham, N., Tyzack, J. D., Ferris, K., & Thornton, J. M. (2017). Mechanism and Catalytic Site Atlas ({M-CSA)}: a database of enzyme reaction mechanisms and active sites. Nucleic Acids Res., 46(D1), D618–D623.

Sankararaman, S., Sha, F., Kirsch, J. F., Jordan, M. I., & Sjölander, K. (2010). Active site prediction using evolutionary and structural information. Bioinformatics, 26(5), 617–624. https://doi.org/10.1093/bioinformatics/btq008

Shoemaker, B. A., Zhang, D., Tyagi, M., Thangudu, R. R., Fong, J. H., Marchler-Bauer, A., … Panchenko, A. R. (2012). {IBIS} (Inferred Biomolecular Interaction Server) reports, predicts and integrates multiple types of conserved interactions for proteins. Nucleic Acids Res., 40(Database issue), D834–40.

Sillitoe, I., Dawson, N., Lewis, T. E. T. E., Das, S., Lees, J. G. J. G., Ashford, P., … Orengo, C. A. C. A. {CATH}: expanding the horizons of structure-based functional annotations for genome sequences, 47 Nucleic Acids Research § (2019). https://doi.org/10.1093/nar/gky1097

Skolnick, J., & Brylinski, M. (2009). FINDSITE: a combined evolution/structure-based approach to protein function prediction. Briefings in Bioinformatics, 10(4), 378–391. https://doi.org/10.1093/bib/bbp017

Sun, J., Wang, J., Xiong, D., Hu, J., & Liu, R. (2016). CRHunter: integrating multifaceted information to predict catalytic residues in enzymes. Scientific Reports, 6(1), 34044. https://doi.org/10.1038/srep34044

Taylor Ringia, E. A., Garrett, J. B., Thoden, J. B., Holden, H. M., Rayment, I., & Gerlt, J. A. (2004). Evolution of enzymatic activity in the enolase superfamily: functional studies of the promiscuous o-succinylbenzoate synthase from Amycolatopsis. Biochemistry, 43(1), 224–229.

Tyzack, J. D., Fernando, L., Ribeiro, A. J. M., Borkakoti, N., & Thornton, J. M. (2018). Ranking Enzyme Structures in the {PDB} by Bound Ligand Similarity to Biological Substrates. Structure, 26(4), 565–571.e3.

Valdar, W. S. J. (2002). Scoring residue conservation. Proteins, 48(2), 227–241.

Wallace, A. C., Borkakoti, N., & Thornton, J. M. (1997). {TESS}: a geometric hashing algorithm for deriving {3D} coordinate templates for searching structural databases. Application to enzyme active sites. Protein Sci., 6(11), 2308–2323.

Wass, M. N., Kelley, L. A., & Sternberg, M. J. E. (2010). {3DLigandSite}: predicting ligand-binding sites using similar structures. Nucleic Acids Res., 38(Web Server issue), W469–73.

Wilkins, A., Erdin, S., Lua, R., & Lichtarge, O. (2012). Evolutionary trace for prediction and redesign of protein functional sites. Methods Mol. Biol., 819, 29–42.

Xue, L. C., Dobbs, D., Bonvin, A. M. J. J., & Honavar, V. (2015). Computational prediction of protein interfaces: A review of data driven methods. FEBS Letters, 589(23), 3516–3526. https://doi.org/10.1016/J.FEBSLET.2015.10.003

Yang, J., Roy, A., & Zhang, Y. (2013). {BioLiP}: a semi-manually curated database for biologically relevant ligand--protein interactions. Nucleic Acids Res., 41(D1), D1096–D1103.

Youn, E., Peters, B., Radivojac, P., & Mooney, S. D. (2007). Evaluation of features for catalytic residue prediction in novel folds. Protein Sci., 16(2), 216–226.

Zhang, T., Zhang, H., Chen, K., Shen, S., Ruan, J., & Kurgan, L. (2008). Accurate sequence-based prediction of catalytic residues. Bioinformatics, 24(20), 2329–2338.

Zhou, N., Jiang, Y., Bergquist, T. R., Lee, A. J., Kacsoh, B. Z., Crocker, A. W., … others. (2019). The CAFA challenge reports improved protein function prediction and new functional annotations for hundreds of genes through experimental screens. Genome Biology, 20(1), 1–23.

