## Supplementary Information for "CATH functional families predict protein functional sites"

### Supplementary Information
### CATH functional families predict protein functional sites

Sayoni Das[1], Harry M. Scholes[2], Christine A. Orengo[2]

[1] PrecisionLife Ltd., Oxford, [2] Institute of Structural and Molecular Biology, University College London, London, UK

#### Section S1. CATH FunFams

##### 1. Generation

Protein domain superfamilies in CATH (Sillitoe et al. 2019) have been sub-classified into functional families, known as FunFams, in which domain relatives are predicted to have highly similar structures and functions. The FunFams are generated by FunFHMMer (Das et al. 2016) which determines an optimal partitioning of the superfamily hierarchical clustering tree on the basis of specificity-determining positions (SDPs) between cluster alignments.

##### 2. Benchmarking

The functional purity of FunFams have been benchmarked using the manually-curated SFLD superfamilies and the EC classification. As a result of their functional purity, FunFams have been shown to perform well in function prediction tasks, such as CAFA competitions(Jiang et al. 2016; Zhou et al. 2019), and using a UniProtKB/Swiss-Prot rollback-assessment of function prediction.

Since FunFams group together protein domains that share similar structure and function, the highly conserved positions in the FunFam alignments tend to be functionally important. These conserved positions can include catalytic sites or allosteric sites for enzymes or binding sites for ligands, metal ions, nucleic acids or interacting with other proteins. The conserved residues in all FunFam alignments have been found to be significantly enriched in known catalytic residues ($p < 1.5 \times 10^{-30}$, Wilcoxon rank sum test) and ligand-binding sites ($p < 7 \times 10^{-22}$, Wilcoxon rank sum test) i.e. FunFams have a greater proportion of catalytic residues and ligand-binding sites within the conserved residues of a domain in comparison to all residues in the domain (Dessailly et al. 2013; Das et al. 2016).

#### Section S2. Supplementary Methods

##### 1. Dataset generation criteria

For each functional site, datasets were generated by:

i. *removing bias and redundancy.* This was done by selecting one domain representative from each FunFam and ensuring that the domains in the set do not have any close function by removing any sequences with >60% sequence identity

ii. *using domains that have at least one known functional site.*

iii. *using domains that can be mapped to a CATH FunFam MSA that have high information content.* An alignment is considered to have high information content if it comprises evolutionarily distant relatives. Information content of MSAs was calculated by measuring the Diversity of Position Score (DOPS) scores generated by Scorecons (Valdar 2002). The DOPS score ranges from 0 to 100 and MSAs with DOPS score of ≥ 70 were considered to have high information content (Das et al. 2016).

All domain definitions used in this analysis were extracted from the CATH v4.2 database (Sillitoe et al. 2019).

#### 2. Datasets generated

**Table S1.** The datasets used for training and testing the FunSite predictors.

| ***Catalytic site (CS) dataset*** *(667 domains)* | | |
|---|---|---|
| | *Training set* | *Hold-out test set* |
| Domains | 565 | 102 |
| CS residues | 1939 | 329 |
| Non-CS residues | 120,723 | 21,095 |
| Ratio of CS/Non-CS residues (approx.) | 1:63 | 1:64 |
| ***Ligand-binding site (LIG) dataset*** | | |
| | *Training set* | *Hold-out test set* |
| Domains | 2026 | 800 |
| LIG residues | 13,870 | 5,634 |
| Non-LIG residues | 310,967 | 103,821 |
| Ratio of LIG/non-LIG residues (approx.) | 1:23 | 1:19 |
| ***Metal-binding site ($LIG_{metal}$) dataset*** | | |
| | *Training set* | *Hold-out test set* |
| Domains | 675 | 240 |
| $LIG_{metal}$ residues | 3,064 | 874 |
| Non-$LIG_{metal}$ residues | 98,697 | 30,015 |
| Ratio of $LIG_{metal}$/Non-$LIG_{metal}$ residues (approx.) | 1:33 | 1:35 |
| ***Protein-protein interaction site (PPI) dataset*** | | |
| | *Training set* | *Hold-out test set* |

| | | |
|---|---|---|
| Domains | 2247 | 599 |
| PPI residues | 78888 | 21028 |
| Non-PPI residues | 287159 | 73813 |
| Ratio of PPI/Non-PPI residues (approx.) | 1:4 | 1:4 |

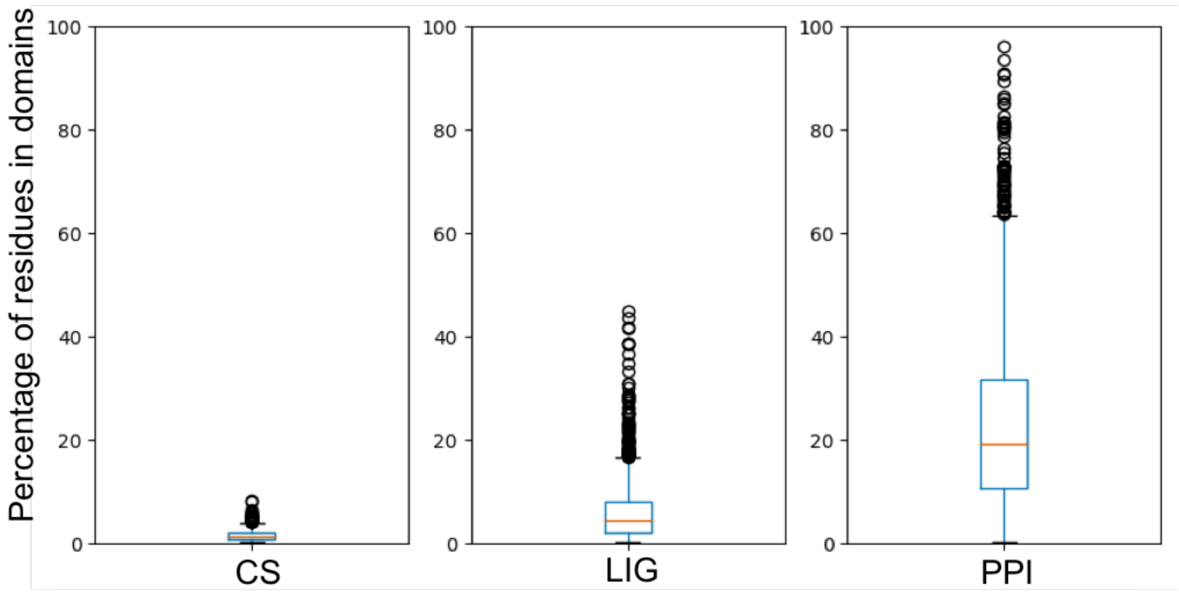


**Figure S1.** Boxplots showing the distribution of known site annotations for all domains in the training dataset for catalytic (CS), ligand-binding (LIG) and protein-protein interaction (PPI) sites.

##### 3. FunSite predictor

**Table S2.** Tree booster parameters in XGBoost that were used to construct the FunSite predictors.

| Parameters | CS | LIG | $LIG_{metal}$ | PPI |
|---|---|---|---|---|
| *n_estimators* | 1000 | 1000 | 1000 | 1000 |
| *learning_rate* | 0.01 | 0.01 | 0.01 | 0.01 |
| *max_depth* | 9 | 6 | 9 | 7 |
| *subsample* | 0.8 | 0.8 | 0.8 | 0.8 |
| *colsample_by_tree* | 1 | 0.8 | 0.8 | 0.8 |
| *gamma* | 0.1 | 1 | 0.1 | 1 |
| *reg_alpha* | 1 | 0.1 | 1 | 1 |
| *scale_pos_weight* | 6 | 6 | 6 | 4 |

#### 4. Features used in the FunSite predictor

##### A. Protein family features

1. *Evolutionary conservation scores*

2. *PSSM and weighted observed percentages (WOP) features*

3. *Conservation scores and predicted functional determinant (FD) scores from Structural Clusters of FunFams*

More information on the protein family features is given in **Section 2.2.2.1.**

##### B. Sequence-based features

1. *Amino acid type:*
   Each residue was encoded by a vector composed of 20 elements, one for each amino acid, where the residue type was set to 1 and the others were set to 0.

2. *Amino acid characteristics*:
   The characteristics of different amino acids were captured using 11 amino acid indices from the AAindex database (Kawashima and Kanehisa 2000). These include bulkiness, charge, hydrophobicity, hydropathicity, hydration potential, hydropathy index, free energy solution, localised electrical effect, mutability, polarity and normalised van der Waals volume.

##### C. Structure-based features

1. *Relative solvent accessibility (RSA)*:
   The relative accessible surface area of total atoms, total side-chain atoms, nonpolar side-chain atoms, polar side-chain atoms, and total main-chain atoms were calculated using the NACCESS program (Hubbard 1992). Residues having RSA of greater than 10% were categorized as surface residues.

2. *Normalized B-factor:*
   The B-factors were extracted from the structure and normalized (per domain) to a value between 0 and 1.

3. *Secondary structure:*
   The secondary structure of each residue was assigned by the DSSP program (Kabsch and Sander 1983).

4. *Cleft or pocket characteristics:*
   The top five largest clefts were predicted in the structure using Speedfill, a modified version of SURFNET (Laskowski 1995). Residues located within a predicted cleft was annotated with the cleft number or 0 otherwise. The depth of the clefts was used as a separate feature. The minimum Euclidean distances of each residue from the three largest pockets were also measured.

5. *Centrality measures:*
   Protein structure networks were created from the structure and three network centrality measures were calculated—degree, betweenness and closeness—using bio3D (Skjærven et al. 2016).

6. *Residue depth and protrusion index:*
   Residue depth and protrusion index were calculated using PSAIA (Mihel et al. 2008).

7. *FOLDX AlaScan:*
   The Gibbs free energies of mutation of each residue to alanine ($DDG_{ala}$) is calculated using the AlaScan command in FoldX (Schymkowitz et al. 2005). Residues with $DDG_{ala}$ > 2 kcal/mol are categorized as putative hotspot residues.

8. *Structural neighborhood characteristics:*
   A Euclidean distance of 5Å was used to identify structural neighbors for each residue. Certain characteristics were then calculated for each residue by taking into consideration the characteristics of its structural neighbors and averaged such as average conservation score, average charge, average hydrophobicity, average number of surface residues, average polarity etc.

#### 5. Features used in the Generic predictors

**Table S3.** List of features in the generic predictors.

| **Type** | **Features** |
|---|---|
| *Protein family features* *(derived from PSI-BLAST)* | Shannon entropy score and other features from PSI-BLAST PSSMs and weighted observed percentages (WOP) matrices, |
| *Sequence Features* | Amino acid residue and their properties<br><br>The amino acid properties include polarity, charge, bulkiness, hydration potential, hydropathy index, hydrophobicity, localized electrical effect, van der Waals vol. normalized. |
| *Structure features* | B-factor,<br><br>Relative Solvent Accessibility,<br><br>Predicted pocket characteristics (number and depth),<br><br>Protein surface characteristics (curvature and depth),<br><br>Secondary structure predictions, |

#### 6. Performance evaluation

The prediction performances are evaluated primarily by the area under the precision-recall curve (PR AUC) and receiver operating characteristic curve (ROC AUC). The PR curve plots precision against recall and the ROC plots TPR against FPR.

The performance of the predictors was also assessed by calculating well-established metrics such as precision, recall (also known as true positive rate or TPR), F1 score, false positive rate (FPR), Matthews correlation coefficient (MCC) and accuracy.

These metrics were calculated as:

$$\text{Precision} = \frac{TP}{TP+FP} \quad (1)$$

$$\text{Recall (TPR)} = \frac{TP}{TP+FN} \quad (2)$$

$$\text{F1 score} = \frac{2\,(Recall\ x\ Precision)}{Recall + Precision} \quad (3)$$

$$\text{FPR} = \frac{FP}{FP+TN} \quad (4)$$

$$\text{MCC} = \frac{TP\ x\ TN - FP\ x\ FN}{\sqrt{(TP+FP)(TP+FN)(TN+FP)(TN+FN)}} \quad (5)$$

$$\text{Accuracy} = \frac{TP+TN}{TP+FN+TN+FP} \quad (6)$$

where TP, TN, FP, and FN denote the numbers of true positives, true negatives, false positives, and false negatives, respectively.

#### Section S3. Supplementary Results

##### 1. New distinguishing features for functional sites

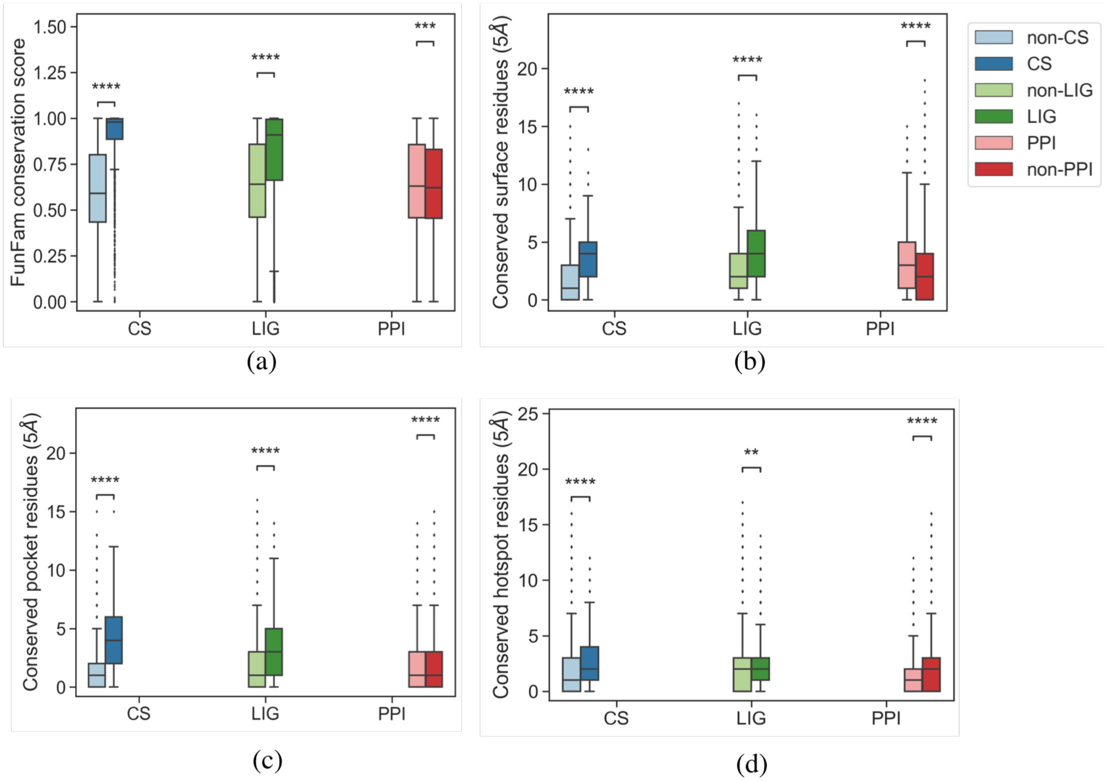


**Figure S2.** A comparison of three FunSite features for CS, LIG and PPI datasets: (a) FunFam conservation score, (b) conserved pocket residue count in structural neighbourhood, (c) average no. of surface residues in structural neighbourhood. Statistical significances were examined by independent t-tests and denoted by asterisks (** $p < 1e\text{-}10$; *** $p < 1e\text{-}20$, **** $p < 1e\text{-}30$).

#### 3. Performance of FunSite predictors

**Table S4.** Performance of FunSite predictors using different metrics with 5-fold cross-validation on the training datasets:

(a) FunSite-CS

| Method | ROC AUC | Precision | Recall | F1 score | MCC | Accuracy |
|---|---|---|---|---|---|---|
| FunSite-CS | 0.951 | 0.718 | 0.762 | 0.739 | 0.696 | 0.924 |
| Generic-CS | 0.938 | 0.628 | 0.769 | 0.691 | 0.639 | 0.903 |

(b) FunSite-LIG

| Method | ROC AUC | Precision | Recall | F1 score | MCC | Accuracy |
|---|---|---|---|---|---|---|
| FunSite-LIG | 0.867 | 0.426 | 0.735 | 0.54 | 0.460 | 0.816 |
| Generic-LIG | 0.846 | 0.386 | 0.726 | 0.505 | 0.418 | 0.791 |

(c) FunSite-PPI

| Method | ROC AUC | Precision | Recall | F1 score | MCC | Accuracy |
|---|---|---|---|---|---|---|
| FunSite-PPI | 0.838 | 0.541 | 0.901 | 0.676 | 0.428 | 0.667 |
| Generic-PPI | 0.785 | 0.493 | 0.921 | 0.642 | 0.355 | 0.605 |

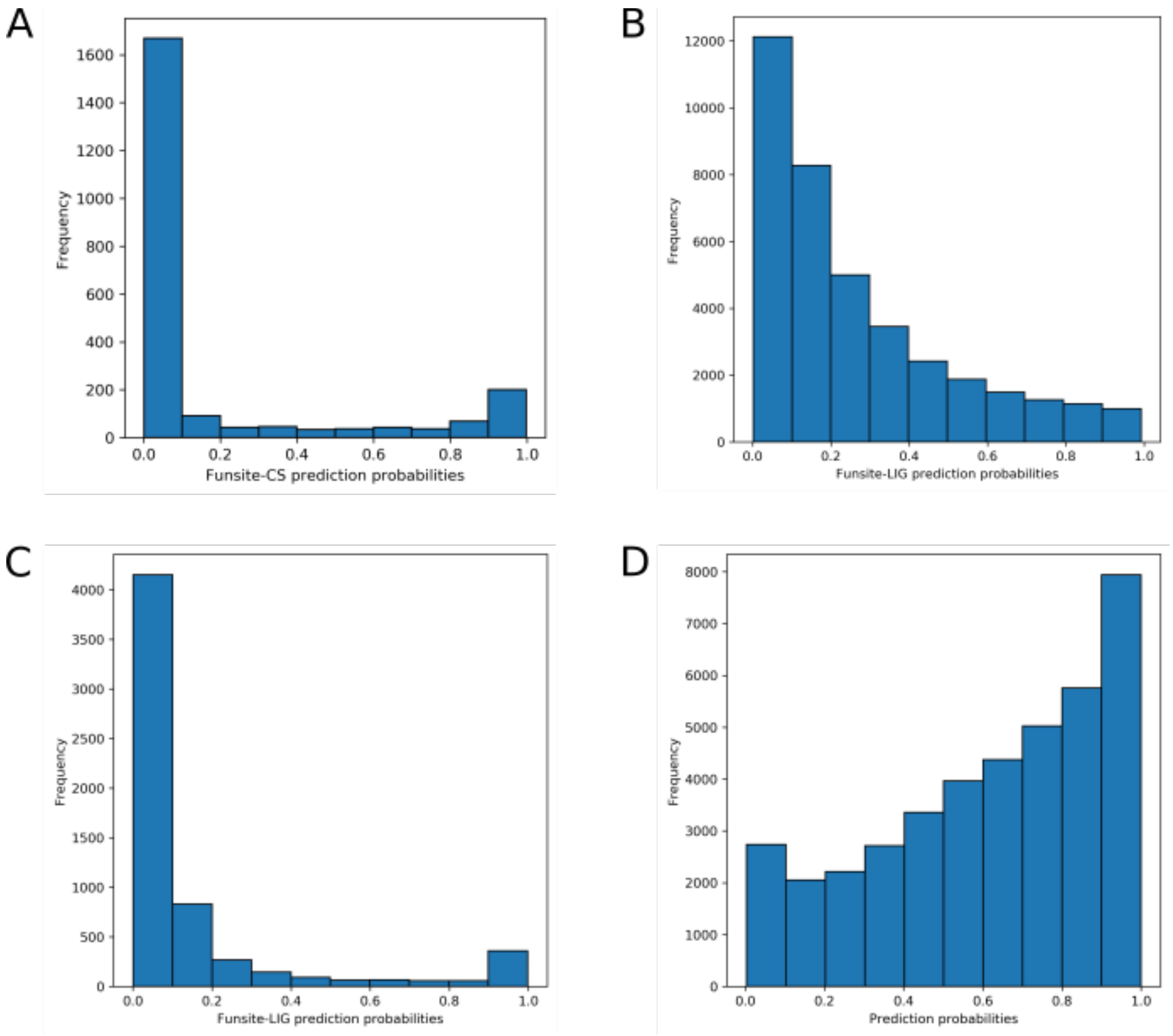


**Figure S3.** Histogram showing prediction probabilities of (a) catalytic sites, (b) ligand-binding sites, (c) metal-binding sites and (d) protein-protein interaction sites generated by FunSite-CS predictor on the hold-out test set.

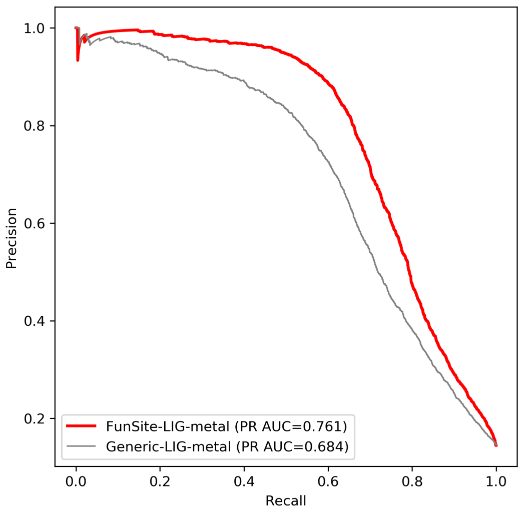


**Figure S4.** Precision-Recall curves showing 5-fold cross-validation results for the FunSite-LIG$_{metal}$ predictor and generic predictor on the training set.

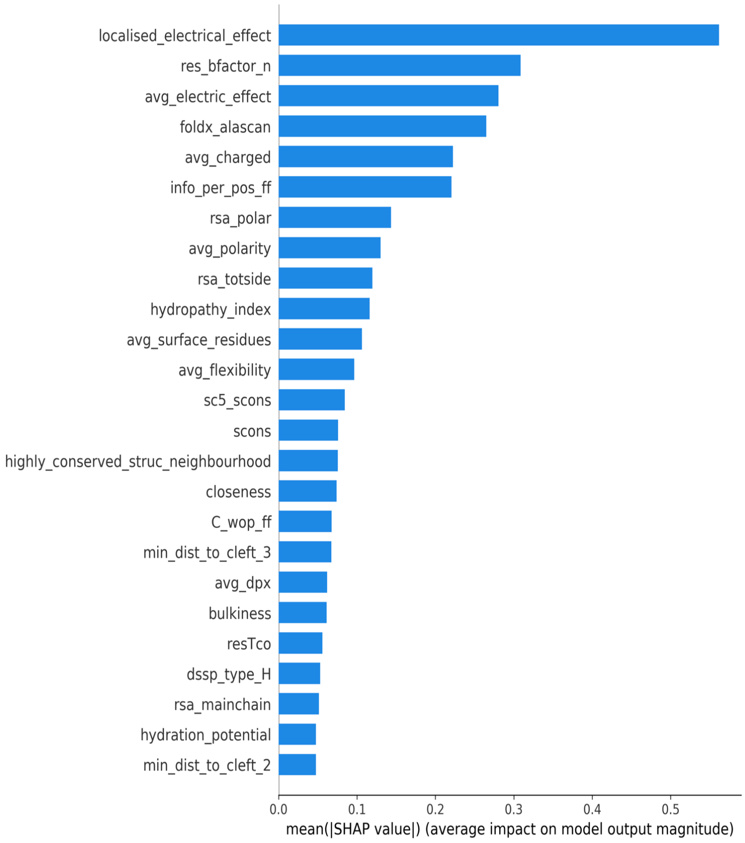


**Figure S5.** SHAP bar plot for $LIG_{metal}$

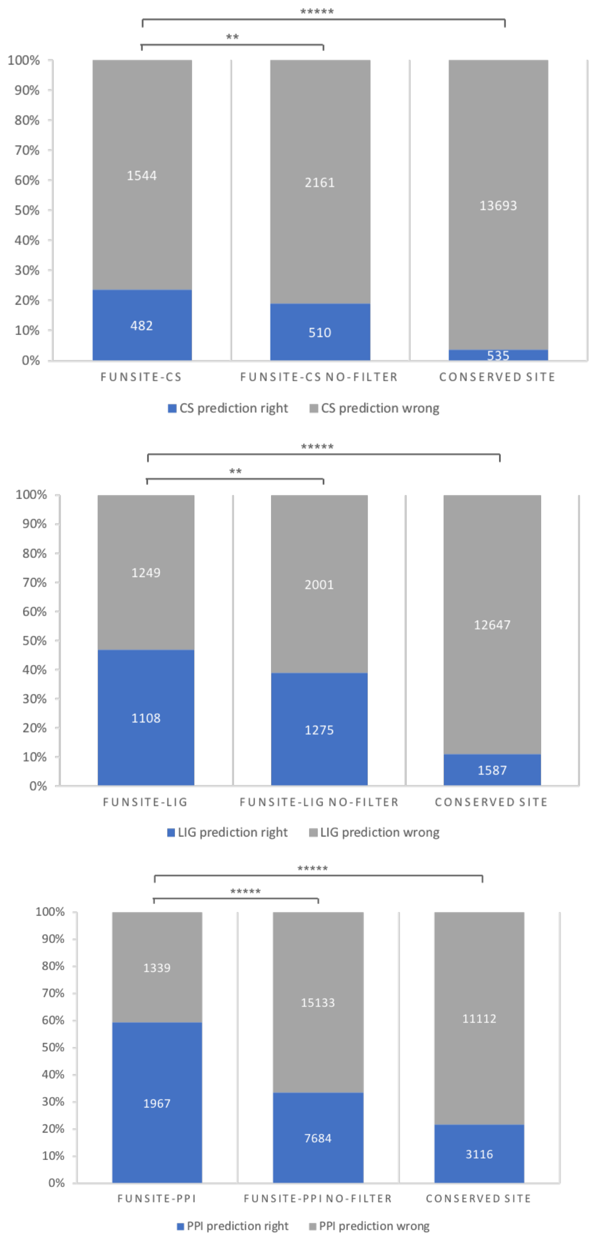


**Figure S6.** The final FunSite predictions (i.e. the top 20 predicted residues) for the CS, LIG and PPI predictors were found to be significantly more enriched in known functional sites compared to the FunSite predictions which hadn't been filtered by removing likely false positives. They were also enriched compared with sets obtained by simply predicting that the conserved sites in FunFams are functional sites.

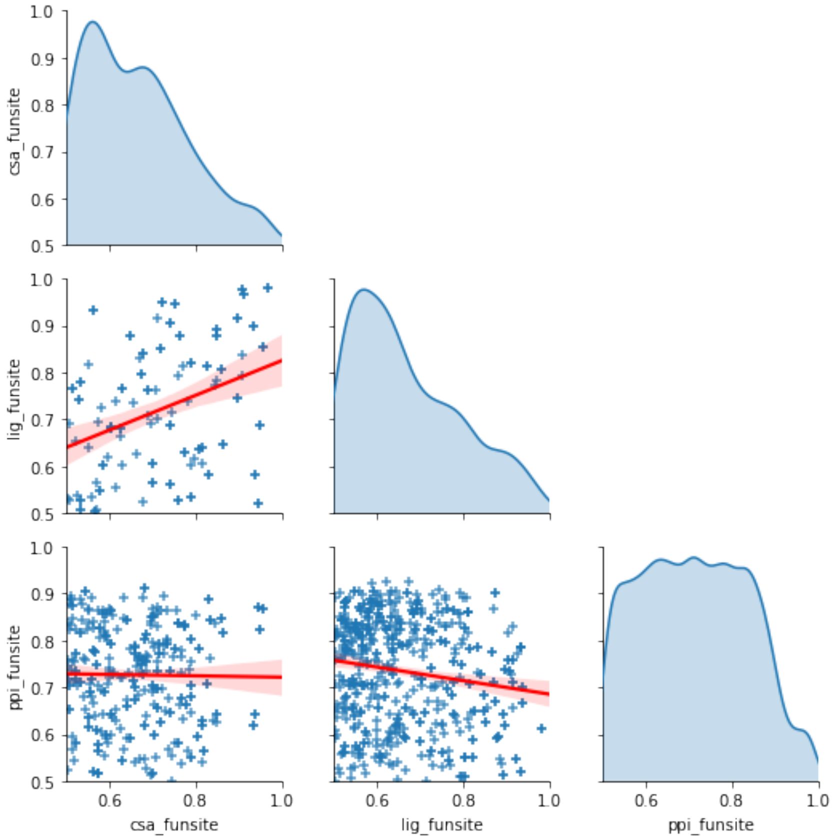


**Figure S7.** Pairwise plots of the prediction scores of catalytic, ligand-binding and protein-protein interaction sites predicted by the full FunSite method for all 175 domains in the combined dataset. Only positively predicted residues for each site are shown.

**References:**


Das, Sayoni, David Lee, Ian Sillitoe, Natalie L Dawson, Jonathan G Lees, and Christine A Orengo. 2016. 'Functional Classification of CATH Superfamilies: A Domain-Based Approach for Protein Function Annotation'. *Bioinformatics* 32 (18): 2889.

Dessailly, Benoit H, Natalie L Dawson, Kenji Mizuguchi, and Christine A Orengo. 2013. 'Functional Site Plasticity in Domain Superfamilies'. *Biochim. Biophys. Acta* 1834 (5): 874–89.

Hubbard, S J. 1992. 'NACCESS: Program for Calculating Accessibilities'. *Department of Biochemistry and Molecular Biology, University College of London*.

Jiang, Yuxiang, T.R. Tal Ronnen Oron, Wyatt T. W.T. Clark, A.R. Asma R. Bankapur, D. D'Andrea, Rosalba Lepore, C.S. Christopher S. Funk, et al. 2016. 'An Expanded Evaluation of Protein Function Prediction Methods Shows an Improvement in Accuracy'. *Genome Biology* 17 (1): 184.

Kabsch, W, and C Sander. 1983. 'DSSP: Definition of Secondary Structure of Proteins given a Set of 3D Coordinates'. *Biopolymers* 22: 2577–2637.

Kawashima, S, and M Kanehisa. 2000. 'AAindex: Amino Acid Index Database'. *Nucleic Acids Res.* 28 (1): 374.

Laskowski, R A. 1995. 'SURFNET: A Program for Visualizing Molecular Surfaces, Cavities, and Intermolecular Interactions'. *J. Mol. Graph.* 13 (5): 307-308,323-330.

Mihel, Josip, Mile Sikić, Sanja Tomić, Branko Jeren, and Kristian Vlahovicek. 2008. 'PSAIA - Protein Structure and Interaction Analyzer'. *BMC Struct. Biol.* 8 (April): 21.

Schymkowitz, Joost, Jesper Borg, Francois Stricher, Robby Nys, Frederic Rousseau, and Luis Serrano. 2005. 'The FoldX Web Server: An Online Force Field'. *Nucleic Acids Res.* 33 (Web Server issue): W382-8.

Sillitoe, Ian, Natalie Dawson, Tony E T.E. Lewis, Sayoni Das, J.G. Jonathan G Lees, Paul Ashford, Adeyelu Tolulope, et al. 2019. *CATH: Expanding the Horizons of Structure-Based Functional Annotations for Genome Sequences*. *Nucleic Acids Research*. Vol. 47.

Skjærven, Lars, Shashank Jariwala, Xin-Qiu Yao, Julien Idé, and Barry J Grant. 2016. 'The Bio3D Project: Interactive Tools for Structural Bioinformatics'. *Biophys. J.* 110 (3): 379a.

Valdar, William S J. 2002. 'Scoring Residue Conservation'. *Proteins* 48: 227–41.

Zhou, Naihui, Yuxiang Jiang, Timothy R Bergquist, Alexandra J Lee, Balint Z Kacsoh, Alex W Crocker, Kimberley A Lewis, et al. 2019. 'The CAFA Challenge Reports Improved Protein Function Prediction and New Functional Annotations for Hundreds of Genes through Experimental Screens'. *Genome Biology* 20 (1): 1–23.
